# Humoral immunogenicity of a Coronavirus Disease 2019 (COVID-19) DNA Vaccine in Rhesus Macaques (*Macaca mulatta*) Delivered using Needle-free Jet Injection

**DOI:** 10.1101/2022.09.12.507647

**Authors:** Alexandra Jay, Steven A Kwilas, Matthew Josleyn, Keersten Ricks, Jay Hooper

## Abstract

A SARS-CoV-2 DNA vaccine targeting the spike protein and delivered by jet injection, nCOV-S(JET), previously shown to protect wild-type and immunosuppressed Syrian hamsters (*Mesocricetus auratus*), was evaluated via two needle-free delivery methods in rhesus macaques (*Macaca mulatta*). The methods included intramuscular delivery of 2 mg per vaccination with the PharmaJet Stratis device and intradermal delivery of 0.4 mg per vaccination with the PharmaJet Tropis device. We hypothesized that the nCOV-S(JET) vaccine would mount detectable neutralizing antibody responses when delivered by needle-free jet injection by either the intradermal or intramuscular route. When delivered intramuscularly, the vaccines elicited neutralizing and variant (Beta, Gamma, and Delta) cross-neutralizing antibodies against SARS-CoV-2 in all six animals after three vaccinations. When delivered at a lower dose by the intradermal route, strong neutralizing antibody responses were only detected in two of six animals. This study confirms that a vaccine previously shown to protect in a hamster model can elicit neutralizing and cross-neutralizing antibodies against SARS-CoV-2 in nonhuman primates. We posit that nCOV-S(JET) has the potential for use as booster vaccine in heterologous vaccination strategies against COVID-19.

## Introduction

Severe Acute Respiratory Syndrome Coronavirus 2 (SARS-CoV-2), the etiologic agent of Coronavirus Disease 2019 (COVID-19), was declared a pandemic by the World Health Organization (WHO) on March 11th, 2020, after its initial discovery in Wuhan, China, in December 2019 [1]. Two years later, nine vaccines have been approved for emergency or full use ¬by at least one stringent regulatory authority and 147 more are in clinical trials [2-4]. Despite a global effort to expeditiously produce safe and efficacious vaccines, supply and manufacturing shortages and availability concerns still exist and still more vaccines are required to meet the world’s demands. Unfortunately, multiple vaccines likely utilizing different platforms will be necessary to accommodate particular difficulties arising in various areas of the world. Two of the three COVID-19 vaccines approved for full use in the United States utilize an mRNA platform (BNT162b2, Pfizer-BioNTech; mRNA-1273, Moderna) and have strict cold chain requirements, which poses significant logistical challenges. Platforms such as DNA vaccines which have a longer shelf life and simpler storage conditions [5] could play a role in future vaccine strategies. Several COVID-19 DNA vaccines are currently in clinical trials [4]. The multi wave nature of the COVID-19 pandemic has made it evident that prolonged periods of insufficient vaccination in a population give the virus ample time to mutate and potentially evade vaccination efforts. Thus, it may become necessary to rapidly develop variant-targeted vaccines to boost both vaccine-generated and natural immunity against the ancestral virus.

Previously we described the development and testing of a needle-free, jet-injected, spike-based SARS-CoV-2 DNA vaccine, nCoV-S(JET), in immunocompetent and immu-nosuppressed Syrian hamster (*Mesocricetus auratus*) models [6]. In that study, neutralizing antibodies levels reached relatively high levels after two vaccinations and significantly reduced disease as measured by body weight loss, lung pathology and virus burden. We were interested in determining if the vaccine would produce similar immunogenicity in a nonhuman primate (NHP) model. We hypothesized that all rhesus macaques (*Macaca mulatta*) would produce detectable neutralizing antibody responses when vaccinated in-tramuscularly (IM) or intradermally (ID) with nCoV-S(JET).

Herein, we describe the testing of nCoV-S(JET) in rhesus macaques using PharmaJet Stratis and Tropis devices. The Stratis and Tropis devices deliver a needle-free jet of liquid intramuscularly or intradermally, respectively. Our primary endpoints for this immunogenicity study were neutralizing antibodies measured by live virus plaque reduction neutralization tests (PRNT) and pseudovirion neutralization assays (PsVNA).

## Materials and Methods

### Virus stocks

A third passage SARS-CoV-2 strain WA-1/2020 viral stock was obtained from the CDC and is from a human non-fatal case isolated in January 2020. A master stock of virus was propagated as described previously [7]. A master stock of virus of NR-54009 (Beta) was also propagated in a similar manner. NR-55674 (Delta) and NR-54984(Gamma) were obtained and used directly from the BEI, American Tissue Culture Collection. All virus work was conducted in BSL-3 containment at USAMRIID.

### Nonhuman Primate Experimental Design

Twelve experimentally-naïve rhesus macaques aged 8 to 15 years of Chinese origin and weighing 5 to 16 kg were utilized for this study. Vaccination groups were randomized and composed of 3 males and 3 females each. Animals underwent a physical examination and bloodwork to confirm lack of underlying disease conditions prior to enrollment on the study. When socially-compatible partners were available, animals were housed in pairs. Animals were sedated with ketamine (Ketamine HCl, 100 mg mL-1, Dechra Veterinary Products, Overland Park, KS; 10 mg kg-1) for all procedures.

Animals received either 0.4 mg of SARS-CoV-2 DNA vaccine, nCOV-S(JET), intra-dermally via PharmaJet Tropis device or 2.0 mg intramuscularly via PharmaJet Stratis de-vice. The immunization volume was split in half and administered on the right and left lower axilla or triceps muscles respectively. All animals were vaccinated on days 0, 21 and 42 and serum and whole blood were collected on days 0, 21, 35, 63, and 168 with blood collection preceding vaccination on days where both procedures occurred. Animals were observed daily for clinical and behavioral abnormalities for the duration of the study.

Research was conducted under an IACUC approved protocol at USAMRIID (USDA Registration Number 51-F-00211728 & OLAW Assurance Number A3473-01) in compliance with the Animal Welfare Act, PHS Policy, and other Federal statutes and regulations relating to animals and experiments involving animals. The facility where this research was conducted is accredited by the AAALAC, International and adheres to principles stated in the *Guide for the Care and Use of Laboratory Animals, National Research Council*, 2011.

### Plaque Reduction Neutralization Test (PRNT)

The PRNTs were conducted as described previously [8]. An equal volume of complete media (MEM containing 10% heat inactivated FBS, 2% Pen/Strep, 100X NEAA, 1% HEPES, 0.1% Gentamycin, and 0.2% Fungizone®) containing SARS-CoV-2 was combined with 2-fold serial dilutions of cMEM containing antibody and incubated at 37°C in a 5% CO2 incubator for 1 hour. Afterwards, the combined virus/antibody mixtures were then added to 6-well plates (180 μl/well) containing 3-day old, ATCC Vero-76 confluent cell monolay-ers and allowed to adsorb for 1 hour in a 37°C, 5% CO2 incubator. Three milliliters of agarose overlay (0.6% SeaKem® ME agarose, 2X EBME with HEPES, 10% heat-inactivated FBS, 100X NEAA, 4% GlutaMAXTM, 2% Pen/Strep, 0.1% Gentamycin and 0.2% Fungizone®) per well was then added and allowed to solidify at room temperature. The plates were placed in a 37°C, 5% CO2 incubator for 2-3 days (variant dependent) and then 2 mL per well of agarose overlay stain (0.6% SeaKem® ME agarose, 2X EBME with HEPES, 5% heat-inactivated FBS, 2% Pen/Strep, 0.1% Gentamycin, 0.2% Fungizone®, 5% Neutral Red) was added. Once plaques could be visualized (one to two days post-staining), plates were removed from the incubator and counted on a light box. PRNT50 titers were determined and are defined as the reciprocal of the highest dilution that results in a 50% reduction in the number of plaques relative to the average number of plaques visualized in the cMEM alone (no antibody) wells.

### Plasmids

The full-length S gene open reading frame, preceded at the N-terminus by the Kozak sequence (ggcacc), was human codon usage-optimized and synthesized by Genewiz (South Plainfield, NJ) and cloned into the Notl-BgIII site of the DNA vaccine vector pWRG for the pWRG/nCoV-S(opt) [6]. The SARS-nCoV-2 S sequence used was the Wuhan coro-navirus 2019 nCoV S gene open reading frame (Genebank accession QHD43416). The plasmids for use in vaccinations were produced commercially and diluted in PBS to 2 mg/mL (Aldevron, Fargo, ND). A second plasmid for the PsVNA was constructed by deletion of 21 amino acids from the COOH terminus of the full length plasmid, pWRG/CoV-S(opt) Δ21 for better in-corporation into pseudovirions [6]. The truncated form was generated using the protocol of 98 °C for 2 min, followed by 25 cycles of, “98 °C for 10 s, 65 °C for 10 s, 72°C for 2 min” and 72°C for 10min. The forward and reverse primer sequences are: Fwd-5′-CCGGCCGCGGCCGCGCCACCATGTTTGTGTTTCTGGTCCTCCTC-3′, Rev-5′-GCTGTTGTAGCTGCGGAAGCTAGTAGTAGGCTAGCAGATCTGCGC-3′. The PCR fragment was gel purified and cloned into the Notl-BgIII site of the DNA vaccine vec-tor pWRG. The pWRG/nCoV-S(opt) plasmid is also called nCOV-S(JET) when combined with jet injection. Additional Δ21 truncated plasmids were created for pseudovirion variant production. The Beta construct was synthesized (CodexDNA, San Diego, CA) directly into the pWRG backbone. For the Delta variant a full-length plasmid was acquired from GenScript (Piscataway, NJ) and then truncated by in-house PCR, and cloned into pWRG using NotI/BamHI. Expression of the spike protein from pWRG/Delta and pWRG/Beta was confirmed by transfection of 293T cells followed by immuno-fluorescence antibody test (IFAT) using heat inactivated (56°C 30 min) human convalescent plasma NRS-53265 (ATCC, Manassas, VA) and compared to empty vector (data not shown).

### Pseudovirion Production

Pseudovirions were produced using the pWRG/CoV-S(opt)Δ21, pWRG/CoV-S(Beta)Δ21, or pWRG/CoV-S(Delta)Δ21 plasmids described above using previously described methods [6]. Briefly, HEK293T cells were transfected with the aforementioned plasmids and after 18 hrs cells were infected with VSVΔG*rLuc. After 3 days at 32°C pseudovirions were concentrated by PEG precipitated and then resuspended in TNE buffer. Pseudovirions were stored at -70°C.

### Pseudovirion Neutralization Assays (PsVNA)

The PsVNA used to detect neutralizing antibodies in sera utilized a nonreplicating vesicular stomatitis (VSV)-based luciferase expressing system [9] with two modifications as previously described [6]. The modifications were (1) no supplemental human complement was added to heat inactivated sera, (2) to reduce any low level residual VSV activity a neutralizing monoclonal antibody was added A regression model (four parameter logistic (4PL) commonly used for bioassays was used to interpolate the PsVNA50 and PsVNA80 titers from which GMTs were calculated.

### MAGPIX Multiplex Immunoassay

MAGPIX using the SARS-CoV 2 spike, S1, RBD, and NP were conducted as previ-ously described [11]. This raw data was converted to a fold change in signal over the pre-bleed by dividing the signal from the study timepoint samples by the pre-bleed sam-ple. Any sample with a fold change greater than four was considered positive.

## Results

Twelve rhesus macaques (*Macaca mulatta)* were vaccinated with nCOV-S(JET) as out-lined in **Table 1**. Sera were collected on days 0, 21, 35, 63, and 168 and evaluated for neu-tralizing antibodies via plaque reduction neutralization (PRNT) and pseudovirion neutralization assays (PsVNA) as well as for binding antibodies via MAGPIX multiplex as-say. Animals were observed daily for signs of adverse reaction, including erythema or other abnormalities at the site of vaccination, changes to food or enrichment consumption, behavioral changes, or change in clinical status. No evidence of adverse reaction following vaccination were noted in any animals during the study.

**Table 1.** Experimental Design. Groups of 6 rhesus macaques each were vaccinated with the nCOV-S(JET) DNA vaccine and sera were collected to evaluate neutralizing and binding antibodies.

| Group | PharmaJet Device | Total Dose per Vaccination | Route | Site | Procedure Schedule (Days) |  |
| --- | --- | --- | --- | --- | --- | --- |
|  |  |  |  |  | Vaccination | Blood Collection |
| ID-DSJI | Tropis | 0.4 mg | ID | Below axilla | 0, 21, 42 | 0, 21, 35, 63, 168 |
| IM-DSJI | Stratis | 2.0 mg | IM | Triceps |  |  |

### Neutralizing antibodies

Pre-vaccination, all animals were negative for neutralizing antibodies by both PRNT and PsVNA (**Figure 1 and 2**). After a single vaccination, two animals in the IM-DSJI group and no animals in the ID-DSJI were positive on at least one of the assays. After two vaccinations, 5 of 6 (83.3%) of the IM-DSJI vaccinated animals developed neutralizing antibodies as compared to 1 of 6 (16.67%) of ID-DSJI vaccinated animals. After 3 vaccinations, 100% of the IM-DSJI group developed neutralizing antibodies as compared to only 33% (2 of 6) of the ID-DSJI group. An additional blood collection was performed on Day 168 to look at durability of the response (**Figure 2**). All but one of the IM-DSJI were still positive in the WA-1 PsVNA, whereas only 50% were still positive by PRNT. The single ID-DSJI vaccinated NHP that was positive by PRNT on Day 63 was still positive on Day 168. That animal, along with two other animals in that group, were positive by WA-1 PsVNA on Day 168. Those two animals had previously been negative by PsVNA on Day 63. Thus, the overall neutralizing antibody positivity rate for the ID-DSJI device was 4 of 6 (66.7%), albeit with low titers.

**Figure 1.**
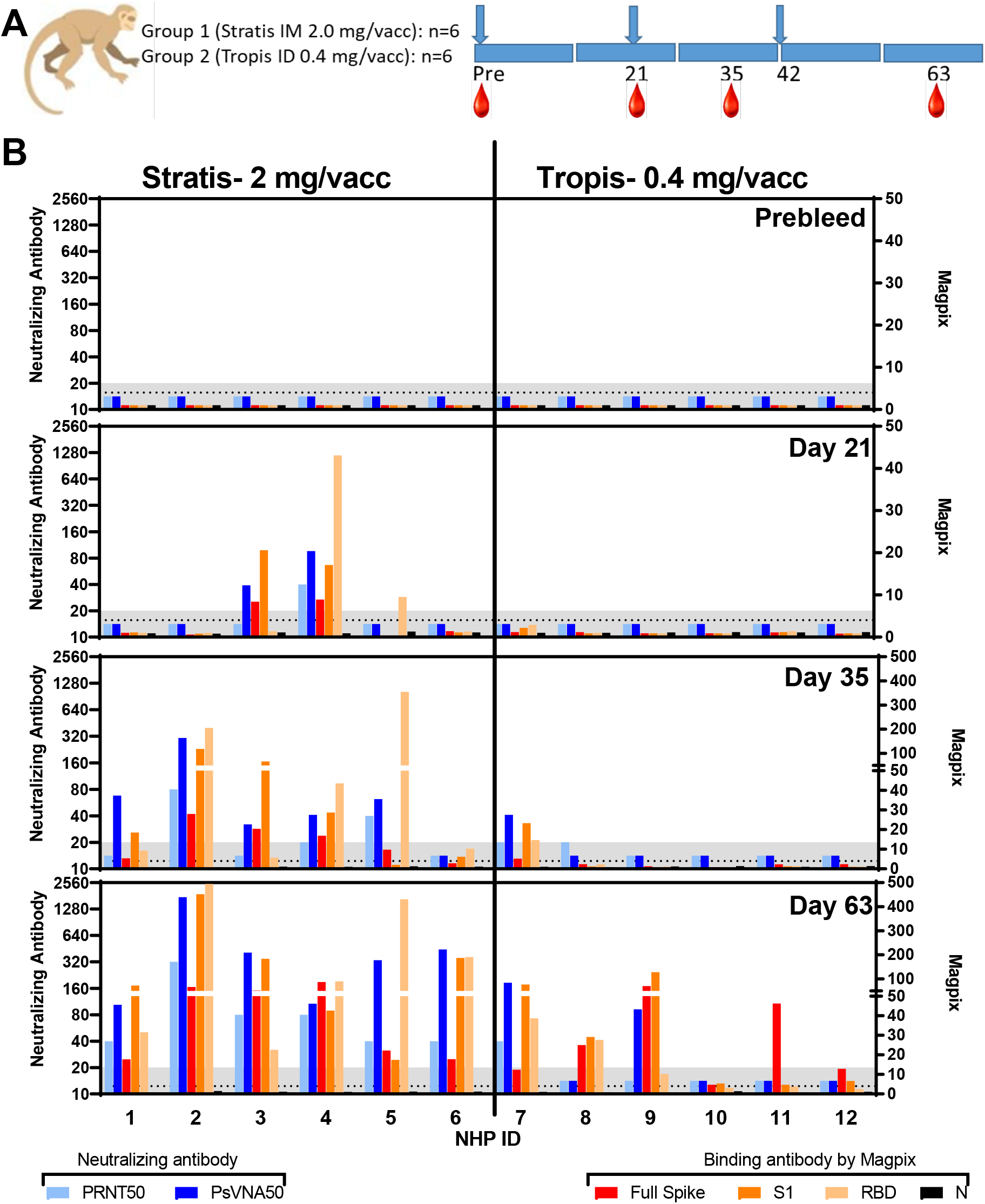
Neutralizing and binding antibody responses. PRNT50, PsVNA50, and Magpix titers from sera collected at various timepoints. **A)** Design. (blue arrows = vaccine dosing; red drops = blood collection time points). **B)** Neutralizing and binding antibody values at the indicated timepoints. Assay lower limits are shown as gray shaded area.

**Figure 2.**
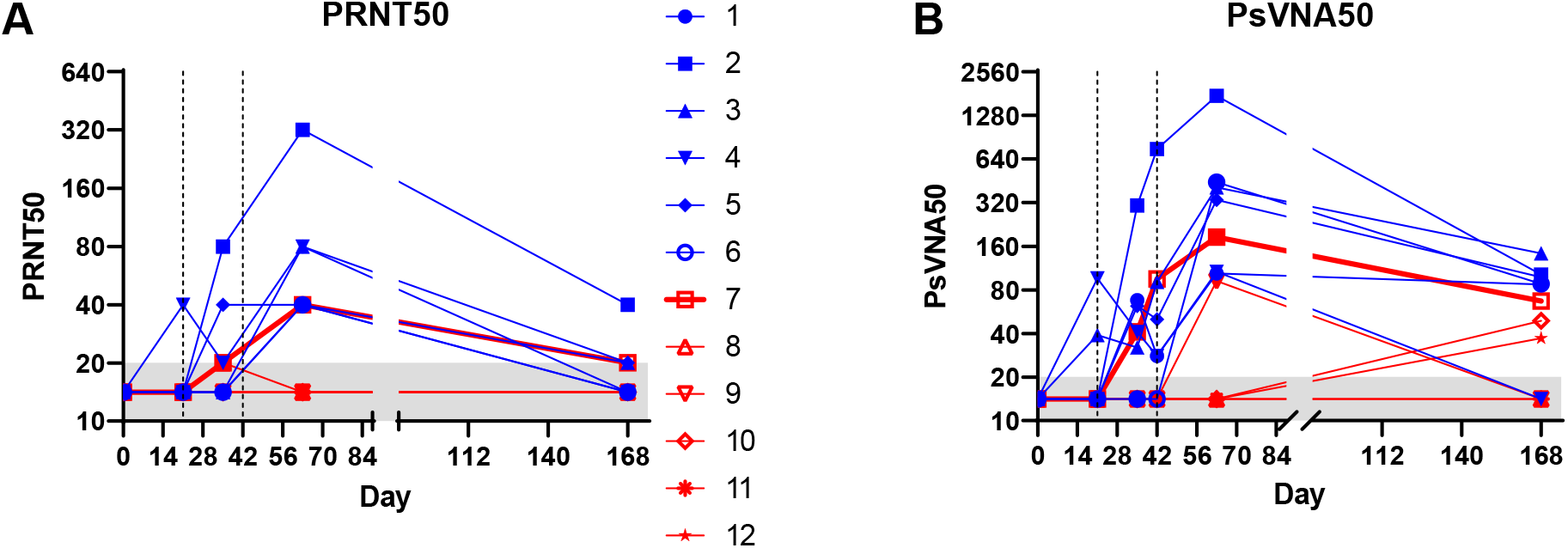
Kinetics of neutralizing antibody responses of individual animals. **A)** PRNT50 and **B)** PsVNA50 titers from sera collected at various timepoints. Vertical dashed lines indicate vaccination timepoints. Blue symbols/lines represent IM-DSJI-vaccinated animals. Red symbols/lines represent ID-DSJI-vaccinated animals. Assay lower limits are shown as gray shaded area.

### Binding antibodies

Magpix assays were performed on sera collected through Day 63 and plotted (**Figure 1**). The assay measured IgG responses against the full spike, S1 domain, receptor binding domain (RBD), and nucleocapsid (N) protein. Predictably, the N re-sponse was negative in all animals at all timepoints. After one IM-DSJI vaccination, the same two animals positive for neutralizing antibodies were also positive for full spike and S1. One of those two animals (#4) also had a high anti-RBD response. A third animal (#5) had detectable anti-RBD antibody but was negative for other proteins. After two vaccina-tions, all of the IM-DSJI animals had at least low levels of binding antibodies against at least two target proteins. Only one animal in the ID-DSJI group had positive binding anti-bodies after two vaccinations (animal #7). After three vaccinations, all of the animals in the IM-DSJI group had improved binding antibody responses and 5 of 6 animals in the ID-DSJI group had detectable binding antibody against at least one target. Animal #10 had almost undetectable responses. The binding data confirmed what was observed in the neutralizing antibody assays: the IM-DSJI device delivering 2 mg was more immunogenic than the ID-DSJI device delivering 5-fold less DNA.

### Cross-neutralizing activity against variants of concern (VOC)

Peak Day 63 sera were evaluated for cross-neutralization activity against Beta, Delta, and Gamma SARS-CoV-2 VOC. Delta and Gamma variants were evaluated by PRNT and Beta and Delta variants by PsVNA. 100% of IM-DSJI vaccinated animals showed cross-neutralization activity against all VOC with the highest reactivity shown against Delta. All ID-DSJI vaccinated animals showed detectable cross-neutralization activity against at least one variant; however only 1 of 6 animals showed cross-neutralization activity against all VOC via both PRNT and PsVNA. Cross-neutralization data are shown in **Figure 3**.

**Figure 3.**
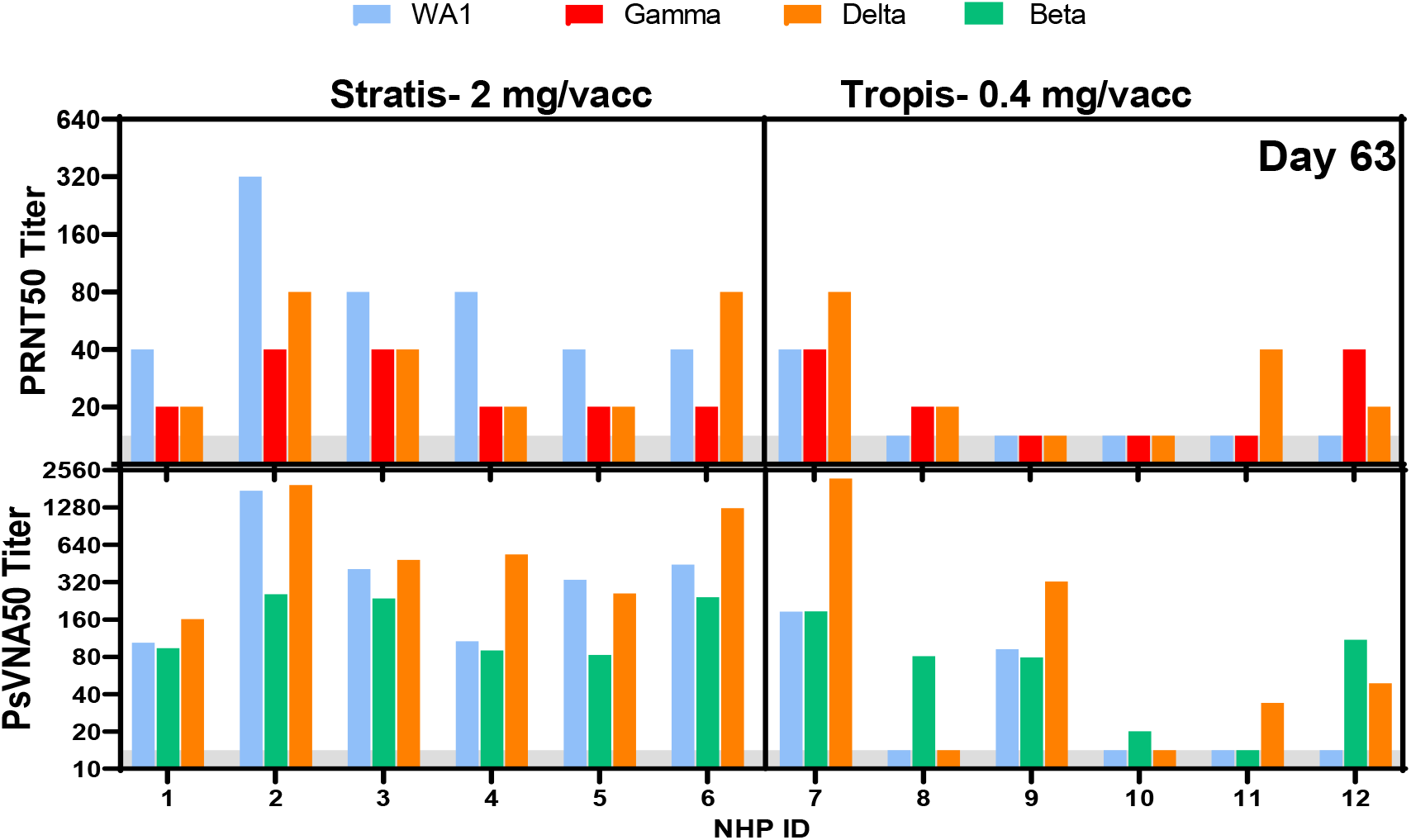
Cross-neutralizing antibody titers against SARS-CoV-2 VOCs from day 63 sera. Assay lower limits are shown as gray shaded area.

## Discussion

This report describes the immunogenicity of the SARs-CoV-2 DNA vaccine nCOV-S(JET) by jet injection in a rhesus macaque model. Our goal was to confirm that the nCOVS(JET) DNA could elicit a detectable neutralizing antibody response and to deter-mine the relative potency of an IM versus ID delivery of the vaccine by jet injection. All IM-DSJI NHPs successfully mounted neutralizing antibodies after 3 vaccinations as com-pared to only 4 of 6 ID-DSJI NHPs. Additionally, neutralizing antibody responses occurred more quickly in the IM-DSJI NHPs, with 100% mounting responses by Day 63 as compared to only 2 of 6 ID-DSJI NHPs. One limitation of this study is that we focused on the humoral immune response and did not conduct a comprehensive assessment of the vaccine-mediated immune response including T cell responses. An additional limitation of this study was that we did not deliver the same dose of DNA by the two different routes. Our rational for delivering different doses was that we delivered the highest dose possible using a standard 2 mg/mL concentration of vaccine drug product. For the IM Stratis device this was 2 mg delivered as two 0.5 mL injections; and for the ID Tropis device this was 0.4 mg delivered as two 0.1 mL injections. Obviously, these doses could be increased by start-ing with more concentrated DNA. Nevertheless, we did determine that the doses used were capable of eliciting neutralizing antibodies in NHPs as measured by both PsVNA and PRNT.

Neutralizing antibody titers failed to rise in the rhesus macaque model as quickly as was seen in Syrian hamsters, with the macaques requiring an additional (second) boost to reach similar antibody titers [6]. The geometric mean titer (GMT) PsVNA50 in hamsters after two vaccinations was approximately 640 whereas in this experiment in NHPs the GMT PsVNA50 was 58 after two vaccinations and 326 after three vaccinations. Similarly, GMT PRNT50 in hamster after two vaccinations was approximately 640 whereas in this experiment in NHPs the GMT PRNT50 was 24 after two vaccinations and 71 after three vaccinations. Using the same PRNT assay we found that macaques infected with WA-1 virus developed PRNT80 titers between 160 and 2560 18 days after exposure [11]. The corresponding PRNT50 titers in that study were between 640 and 5120 (data not shown). It is possible that the NHPs received an inadequate dose as compared to the hamsters when juxtaposed on a per weight basis. Hamsters received a total of 0.4 mg nCOV-S(JET) intramuscularly over three vaccinations, as compared to the 6.0 mg received intramuscu-larly in the NHPs. While our NHP dosing is similar to other vaccines utilizing DNA plat-forms, the dose used in our small animal model was substantially higher [12-14]. Since our previous work in hamsters only explored vaccine efficacy via the intramuscular route we cannot truly compare potential discrepancies in the dosing per weight, but DNA vaccines historically have demonstrated the highest immunogenicity when administered intramuscularly as compared to other routes [15]. Additionally, we are unable to evaluate the protective efficacy of our vaccine without having challenged the animals with SARS-CoV-2.

A stabilized spike DNA vaccine delivered by needle injection protected NHPs from disease [16]. The neutralizing antibody titers in those protected animals as measured by a PsVNA was just over 100. Similarly, a spike-based DNA vaccine termed ZyCoV-D delivered three times to rabbits intradermally using the Tropis device elicited neutralizing an-tibody titer of 108 in a microneutralization test [14]. The same vaccine administered 2 mg/vaccination three times using Tropis to humans in a phase 1 clinical resulted in a neutralizing antibody GMT of 39.17 [17]. Thus, the neutralizing antibody we observed, at least in the IM-DSJI group, were comparable to titers that were protective in NHPs and similar to the titers elicited in rabbits of a vaccine that has been approved in India.

The nCOV-S(JET) vaccine delivered using the IM Stratis device exhibited impressive VOC cross-neutralizing activity, especially when measured by the PsVNA. All of the animals had PsVNA50 titers against WA-1, Beta, and Delta of at least 80 and were as high as 1280. Interestingly, the responses against Delta appeared to as high, or higher, than against WA-1. The PRNT50 were less impressive, but still all of the IM-DSJI NHPs had PRNT50 titers of at least 20 and has high as 80 against the Gamma and Delta VOC. Un-surprisingly, the low dose of DNA delivered by ID Tropis device had lower cross-neutralizing responses. Only two animals (#7 and #9) had detectable cross-neutralizing antibodies against all VOC tested (by PsVNA) and only one of those animals was also positive against all VOC tested by PRNT (animal #7). These two animals also had the most robust binding response as measured by Magpix indicating high levels of binding antibodies is predictive of a potent neutralizing and cross-neutralizing re-sponse.

At the doses used in this study, two boosts were necessary to produce responses that would likely be beneficial to the vaccine recipient (i.e., neutralizing antibody responses). This number of vaccinations is not well suited for rapid, large scale vaccination cam-paigns, but may be most appropriate for heterologous boosting strategies. Heterologous boosting, wherein the booster vaccine is of a different platform than that used to complete the primary vaccination series, allows for increased flexibility, a significant advantage in the current landscape of vaccine shortage, and has the potential for reduced reactogenicity and increased immunogenicity. In a clinical trial utilizing 3 COVID-19 vaccines (Moderna mRNA-1273; Janssen Ad26.COV2.S; Pfizer-BioNTech BNT162b2) authorized by the U.S. Food and Drug Administration (FDA), fully vaccinated adults receiving heterologous boosts displayed similar reactogenicity and increased immunogenicity as measured by neutralizing and binding antibody titers as compared to homologous boosts for all combinations [18]. Similar evidence exists for heterologous priming strategies, wherein the second vaccination dose utilizes a different platform than that of the first, as well. A systematic review found heterologous priming with BNT162b2 in participants partially vaccinated with ChAdOx-1 produced robust immunogenicity and tolerable reactogenicity [19]. With promising initial findings, heterologous vaccination strategies warrant further research into long-term safety profiles as well as identification of optimal combinations and dosing regimens. As of October 21, 2021, the FDA authorized use of three aforemen-tioned COVID-19 vaccines for use as heterologous boosts [20]. At present, the WHO has yet to recommend heterologous boosts as a general recommendation.

Nucleic acid vaccines have the potential to be highly amenable to use where rapid response and rapid adaptation is needed. This has clearly been demonstrated by the mRNA COVID vaccines that were rapidly advanced from discover to licensure. The DNA vaccine used in this study was not formulated with lipid nanoparticles and did not re-quire any adjuvant or electroporation.

The vaccine was delivered by a technology that does not require electricity and uses relatively inexpensive disposable needle-less syringes. These devices developed by PharmaJet are FDA 510-k cleared and the Tropis device is currently utilized by Zydus Cadila Healthcare for their three dose COVID DNA vaccine that has been approved for emergency use in India. Our data confirm the immunogenicity of simple nCoV-SARS-2 DNA delivered by jet injection. For this particular DNA vaccine, increases in dose or other steps to increase potency are warranted for a standalone vac-cine; however, it is possible that the current configuration could be used to rapidly boost existing immunity using DNA vaccines encoding VOC such as Omicron.

## Acknowledgements

We would like to thank Kevin Nemelka, Cara Reiter, Brett Taylor, Joseph Royal, Philip Bowling, Brianna Marion, and the rest of the Veterinary Medicine leadership for their support during this study as well as the In Vivo Studies veterinary technicians for their outstanding technical support and all of the Veterinary Medicine Division animal caretakers for their dedicated care. We would also like to thank Tamara Clements of the Diagnostic Systems Division for conducting the Magpix immunoassay. Opinions, interpretations, conclusions, and recommendations are those of the author and are not necessarily endorsed by the U.S. Army.

